# Actomyosin contractility modulates Wnt signaling through adherens junction stability

**DOI:** 10.1101/220178

**Authors:** Eric T. Hall, Elizabeth Hoesing, Endre Sinkovics, Esther M. Verheyen

**Affiliations:** Department of Molecular Biology and Biochemistry, Centre for Cell Biology, Development and Disease, Simon Fraser University, Burnaby, V5A 1S6, Canada

## Abstract

Mechanical forces can influence the canonical Wnt signaling pathway in processes like mesoderm differentiation and tissue stiffness during tumorigenesis, but a molecular mechanism involving both in a developing epithelium and its homeostasis is lacking. We identified that increased non-muscle myosin II activation and cellular contraction inhibited Wnt target gene transcription in developing *Drosophila*. Genetic interactions studies identified this effect was due to myosin-induced accumulation of cortical F-actin resulting in clustering and accumulation of E-cadherin to the adherens junctions. E-cadherin titrates any available β-catenin, the Wnt pathway transcriptional co-activator, to the adherens junctions in order to maintain cell-cell adhesion under contraction. We show that decreased levels of cytoplasmic β-catenin result in insufficient nuclear translocation for full Wnt target gene transcription. Our work elucidates a mechanism in which the dynamic activation of actomyosin contractility refines patterning of Wnt transcription during development and maintenance of epithelial tissue in organisms.

## Introduction

The Wnt signaling pathway [Wingless (Wg) in *Drosophila*], is highly conserved across metazoans and essential during development and tissue homeostasis for the regulation of proliferation and patterning [1]. Wnt signaling achieves proper biological outcomes through extensive crosstalk with other signaling pathways. Recent studies have begun to elucidate how mechanical forces may also play critical roles in regulating signaling pathways during development [2]. Here, we identified a molecular mechanism in which actomyosin activation and the resulting contractile forces within a cell can regulate Wnt signaling.

The canonical Wnt/Wg pathway centers on the stabilization and localization of the key effector protein, β-catenin (β-cat) [Armadillo (Arm) in *Drosophila*]. β-cat is continuously produced in most cells for its roles in both the formation and maintenance of adherens junctions (AJs) and as a transcriptional activator for Wnt signaling [3]. AJs are major epithelial cell-cell adhesion complexes that maintain tissue integrity in response to external forces like morphogenesis [4]. AJs mainly form an apical-lateral belt-like structure around cells, holding neighbouring cells together through the homophilic binding of the transmembrane protein E-cadherin (E-cad). β-cat binds to the cytoplasmic tail of E-cad and to α-catenin, which interacts with the actin cytoskeleton. Thus AJs act as mechanical force integration sites across cells and at a tissue level [5].

In the absence of a Wnt ligand, cytoplasmic β-cat is targeted for degradation by a multi-protein destruction complex which assembles on the scaffolding protein Axin and includes kinases that phosphorylate β-cat, targeting it for ubiquitination and subsequent proteasomal digestion. Upon Wnt/Wg binding to its coreceptors Frizzled (Fz) and LRP/Arrow, Dishevelled (Dvl/Dsh) is recruited to Fz and triggers the recruitment of the destruction complex to the membrane. This event disrupts the destruction complex, allowing β-cat to accumulate and translocate to the nucleus, where it acts with T-cell factor (TCF)/lymphoid enhancer factor (LEF) transcription factors to initiate target gene expression [6]. Disruptions of the core components or regulatory proteins have been found in numerous cancers and developmental disorders [1].

Recent studies suggest that canonical Wnt signaling and mechanical forces are integrated in regulation of development and homeostasis. Wnt activation can drive mechanical strain-induced cell proliferation [7] as well as activation of non-muscle myosin II (NMII) leading to morphogenesis [8]. Conversely, force induction and subsequent cytoskeletal rearrangements can regulate Wnt signaling, but these studies have typically focussed on extracellular matrix (ECM) stiffness in stem cells or on tumorigenic situations [9–12]. Insight is lacking on the role of force induction and its regulation of Wnt activation in normal developing epithelial tissues. Recently multiple components of myosin phosphatase were identified in a kinome and phosphatome RNA interference (RNAi) screen to identify novel phospho-regulators of Wnt signaling in developing *Drosophila* larvae [13].

Myosin phosphatase is the major inhibitor of NMII in cells. It consists of two major proteins, either one of two targeting subunits, the myosin phosphatase targeting protein MYPT1/2 (Myosin binding subunit (MBS) in *Drosophila*) or MYPT3 (*Drosophila* Mypt-75D) and the catalytic protein phosphatase type 1β (PP1β) subunit (encoded by *flapwing (flw)* in *Drosophila*) [14]. Myosin phosphatase inactivates NMII by dephosphorylating Thr-18 and Ser-19 (*Drosophila* Thr-20 and Ser-21), the two critical activation residues of the regulatory light chain (encoded by *spaghetti squash* (*sqh*) in *Drosophila*) of NMII [15,16].

NMII is the major actin-binding motor protein that drives actomyosin cytoskeletal contraction. Its activation controls a diverse range of mechanisms included cell shape, adhesion, migration, cell cycle and cell division [17]. NMII regulatory light chain phosphorylation and the resulting contractile force activity can be induced by numerous kinases, some of which also phosphorylate and inhibit myosin phosphatase [17]. Several upstream Rho GTPases that activate myosin kinases and thus stimulate NMII can inhibit Wg activity in *Drosophila*, but a fully defined mechanism is not known [18]. Here we show that actomyosin-based force generated by NMII stimulation within and across cells in an epithelium can modulate Wnt signaling and tissue patterning by preferentially stabilizing cell-cell adhesion at the AJs to maintain tissue integrity at the expense of transcription and patterning.

## Results

### Myosin Phosphatase promotes activation of Wg signaling

Components of myosin phosphatase were identified in an RNAi screen due to their ability to modulate Wg target gene expression in the wing imaginal disc [13]. The Wg target gene *Distal-less (Dll)* is expressed in a broad domain within the wing pouch (Fig 1A) [19]. Expression of *mypt-75D-RNAi* or *flw-RNAi* in the posterior domain of the wing imaginal disc using *hedgehog (hh)-Gal4* (referred to as *hh>mypt-RNAi)* caused a strong reduction in *Dll* transcription (Figs. 1A’, S1A). Adult flies had a dramatic size reduction in the posterior of the wing blade as well as notches and loss of wing bristles, hallmarks of reduced Wg signaling (Fig 1E). The Wg ligand is expressed in a band 2-3 cells wide along the dorsoventral (D/V) boundary (Fig 1B), which was unaffected in GFP-marked actin flip-out clones expressing *mypt-75D-RNAi or flw-RNAi* (Figs. 1B’, S1B), indicating that reduced myosin phosphatase was not disrupting ligand production to inhibit Wg signaling.

**Fig 1.**
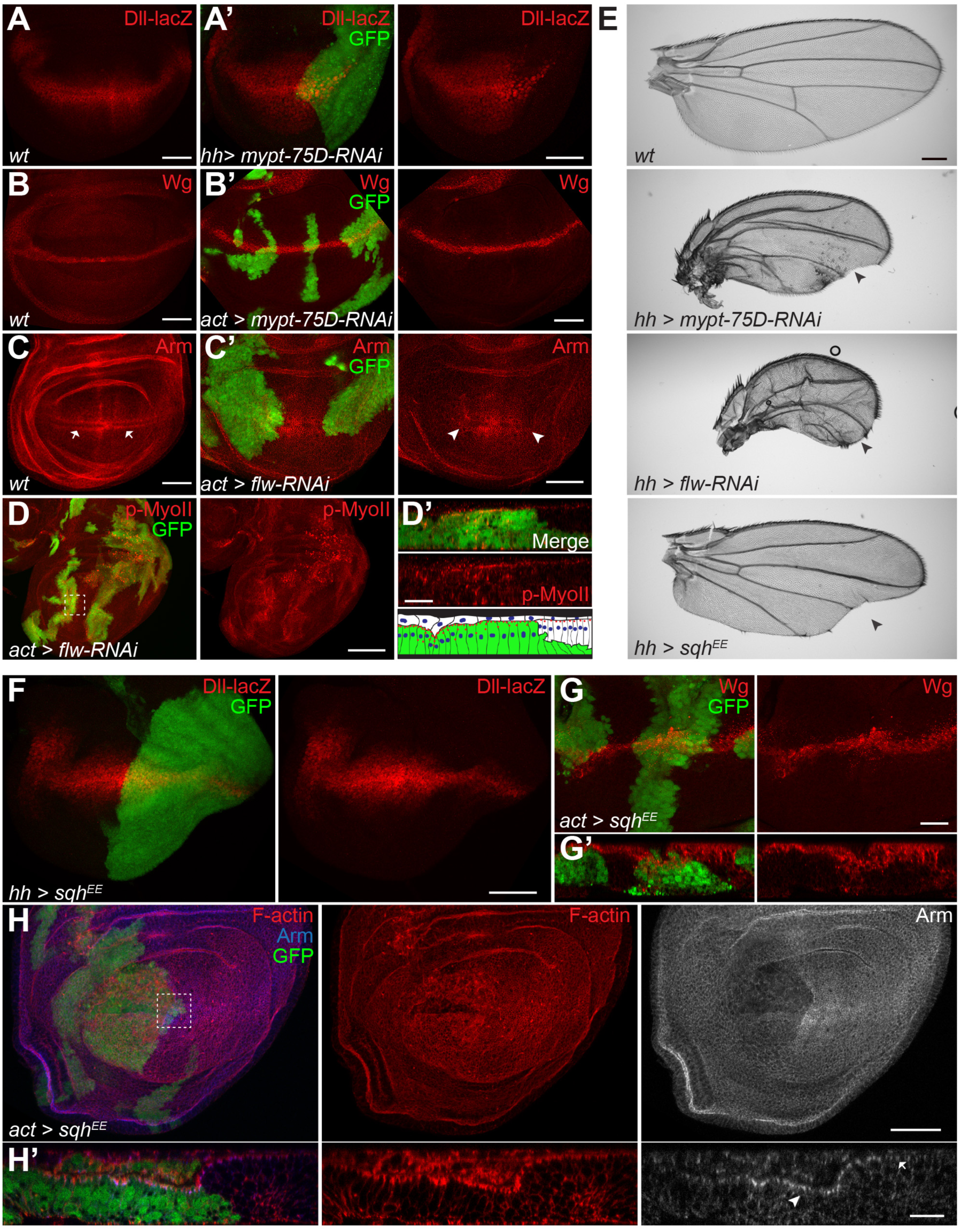
Myosin phosphatase and NMII regulate Wg activity during wing development. (A,A’) *Dll-lacZ* expression in wild type (A) and *hh-Gal4* driving *gfp* and *mypt-75D-RNAi* (A’) third-instar wing imaginal discs. (B,B’) Wg protein expression in wild type (B) and GFP-marked actin flip-out clones driving *mypt-75D-RNAi* (B’). (C,C) Arm stabilization pattern in wild type (C, arrows) and in flip-out clones driving *flw-RNAi* (C’, arrowheads). (D,D’) p-MyoII stained in *flw-RNAi* flip-out clones. Cross section seen in (D’) is the magnified dashed line area of (D). (E) Adult wings of wild type, and *hh-Gal4* driving *mypt-75D-RNAi, flw-RNAi,* or *sqh^EE^* (arrowheads mark loss of bristles and wing margins). (F) *hh>sqh^EE^, gfp* stained *for Dll-lacZ* expression. (G, G’) Total Wg in *sqh^EE^* flip-out clones, and cross section showing cell constriction. (H,H’) GFP flip-out clones driving *sqh^EE^* stained for F-actin and Arm. (H’) Cross section (magnified dashed line area of (H)) shows apical F-actin and Arm (H’ arrowhead vs. arrow). Scale bars: (A-C,F,H) 50 μm, (D) 100 μm, (D’,G,G’,H’) 20 μm, (E) 300 μm.

We next examined the stability of the key effector, Arm, which accumulates at the highest concentrations in the cytoplasm and nucleus in two visible bands of cells flanking the Wg-producing cells (Fig 1C arrows) [20]. Flip-out clones expressing*flw-RNAi* (Fig 1C’) or *hh* > *mypt-75D-RNAi* (Fig S1C) both caused reduced stabilized Arm, seen by the loss of bands.

We confirmed that the reduction of Arm was not due to cell death, as staining for the apoptotic marker cleaved caspase-3 (C.Casp-3) was not elevated following knockdown *of flw* or *mypt-75D* (Fig S1B,C). Knockdown of the other targeting subunit, MBS, gave similar results to *flw-RNAi* or *mypt-RNAi,* but could induce cell death, and was therefore not used in further experiments (Fig S2). Taken together these results demonstrate a previously uncharacterized role for myosin phosphatase in the promotion of Wg signaling in *Drosophila.*

### Increased NMII activity inhibits Wg signal activation

The key role of myosin phosphatase is to dephosphorylate and inactivate NMII. We confirmed that knockdown of *flw* or *mypt-75D* lead to hyperactive phospho-NMII. RNAi clones of either *flw* or *mypt-75D* had increased phosphorylated Sqh (p-MyoII) (Figs. 1D, S1D). Cross sections showed other phenotypes associated with elevated NMII activation (Fig 1D’). The wing imaginal disc consists of tightly packed columnar epithelial cells where Wg signaling occurs, and a thin layer of squamous peripodial epithelium above its apical surface (Fig 1D’ cartoon) [21]. *flw-RNAi* cells had elevated levels of p-MyoII, were constricted and formed clefts (Fig 1D’) and had reduced apical surface area, indicated by E-cad::GFP (Fig S1E), another sign of increased NMII activity, and a proxy to force generation [22].

We next asked if directly activating NMII could phenocopy the loss of Wg signaling seen with myosin phosphatase knockdown. An activated phosphomimetic NMII regulatory light chain (Sqh^EE^) could inhibit Wg target gene expression and induce notched wings in adults (Fig 1E,F). Like myosin phosphatase knockdown, Sqh^EE^ could induce tissue constriction (Fig 1G’,H’), but did not affect Wg protein levels (Fig 1G). Cells with increased NMII activity also had elevated levels of F-actin (Fig 1H,H’). Activated NMII binds actin and stabilizes filaments as it pulls them together, reducing their turnover rate and causing an overall increase in F-actin in the cell [23]. Like reduced myosin phosphatase, Sqh^EE^ could reduce levels of stabilized cytoplasmic and nuclear Arm (Fig 1H), but cross sections showed increased levels of AJ Arm at the apical surface of constricting cells (Fig 1H’). This suggested that a defect in Arm distribution within cells having increased NMII activity may underlie the reduction in Wg targets. Moreover, we found that directly increasing myosin activation can modulate Wg activation. We next wanted to ask what happened if we did the inverse experiments in which myosin activity was reduced.

Overexpression of Mypt-75D (*hh>mypt-75D,* inactivating NMII) surprisingly caused a dramatic loss of *Dll* expression in the posterior of the wing disc (Fig S3B’, D). This is most likely due to elevated cell death as shown with elevated levels of cleaved caspase 3 (Fig S3B-B’’). Confirming this, upon co-expression of the baculoviral P35 anti-apoptosis protein, *Dll* transcription was restored (Fig S3C-C’’)[24], suggesting loss of Wg activity from overexpression of Mypt-75D was indirect and due to cell death. Interestingly co-expression of activated Sqh^EE^ could also partially restore *Dll* expression in a Mypt-75D overexpression background (Fig SD-F), indicating that inactivating NMII caused reduced cell viability, which could be partially restored by ectopic activated NMII. Furthermore, even cells mutant for functional NMII expressing P35, were non-viable and quickly extruded from the tissue apically or basally (Fig S3H, arrows).

Consistent with the overexpression studies with myosin phosphatase, we found that the knockdown of total NMII (*hh>sqh-RNAi)* predominantly induced widespread cell and organismal death, and the few wing discs that survived exhibited a dramatic reduction in total area of the posterior domain of the wing disc and a complete loss of *Dll* expression (marked by the dotted line, Fig S3I). Knockdown of myosin phosphatase components in this background could partially restore *Dll* expression and some of the growth defects (Fig S3J-M), suggesting that low levels of NMII affect cell viability, but that activating the remaining NMII complexes (by removal of myosin phosphatase) can partially restore viability, and as a consequence, Wg activity. However, tissue with severely reduced NMII frequently becomes apoptotic and becomes extruded from the surrounding epithelium, making it exceedingly difficult to analyze. Although this may be a distinct mechanism for possibly regulating Wg activity, the widespread induction of cell death and extrusion due to loss of NMII will not be further characterized in this study.

### NMII activation reduces nuclear Arm independently of the destruction complex

As our results indicate myosin phosphatase likely affects Wg signaling through inactivation of NMII (Fig S3N), in subsequent experiments knockdown of myosin phosphatase will be used as a proxy for specifically stimulating NMII. Since increased NMII activity led to reduced stabilized Arm and a loss of Wg target gene expression, we next asked if NMII could affect destruction complex proteins in *Drosophila* salivary gland cells, as their large cells are ideal for studying protein localization *in vivo,* and glands have been characterized with respect to Wg signaling [25].

Knockdown of *flw* using *dpp-Gal4* resulted in a loss of the Wg target gene *fz3* (Fig 2A), indicating that activated NMII can also inhibit Wg signaling in the salivary gland. After Wg binds to its receptors in the proximal cells of the salivary gland, Dsh, Axin and other the components of the destruction complex are recruited to Fz (Fig 2B), causing inactivation of the complex [26]. *flw-RNAi* had no effect on the cell surface distribution of Dsh-GFP or FLAG-Axin (Fig 2B), compared to other proteins previously identified to affect recruitment to the membrane [25], indicating that Wg’s regulation of the destruction complex still occurs in cells with elevated NMII activity. These results suggest that increased NMII activity affects Wg signaling downstream of the receptors.

**Fig 2.**
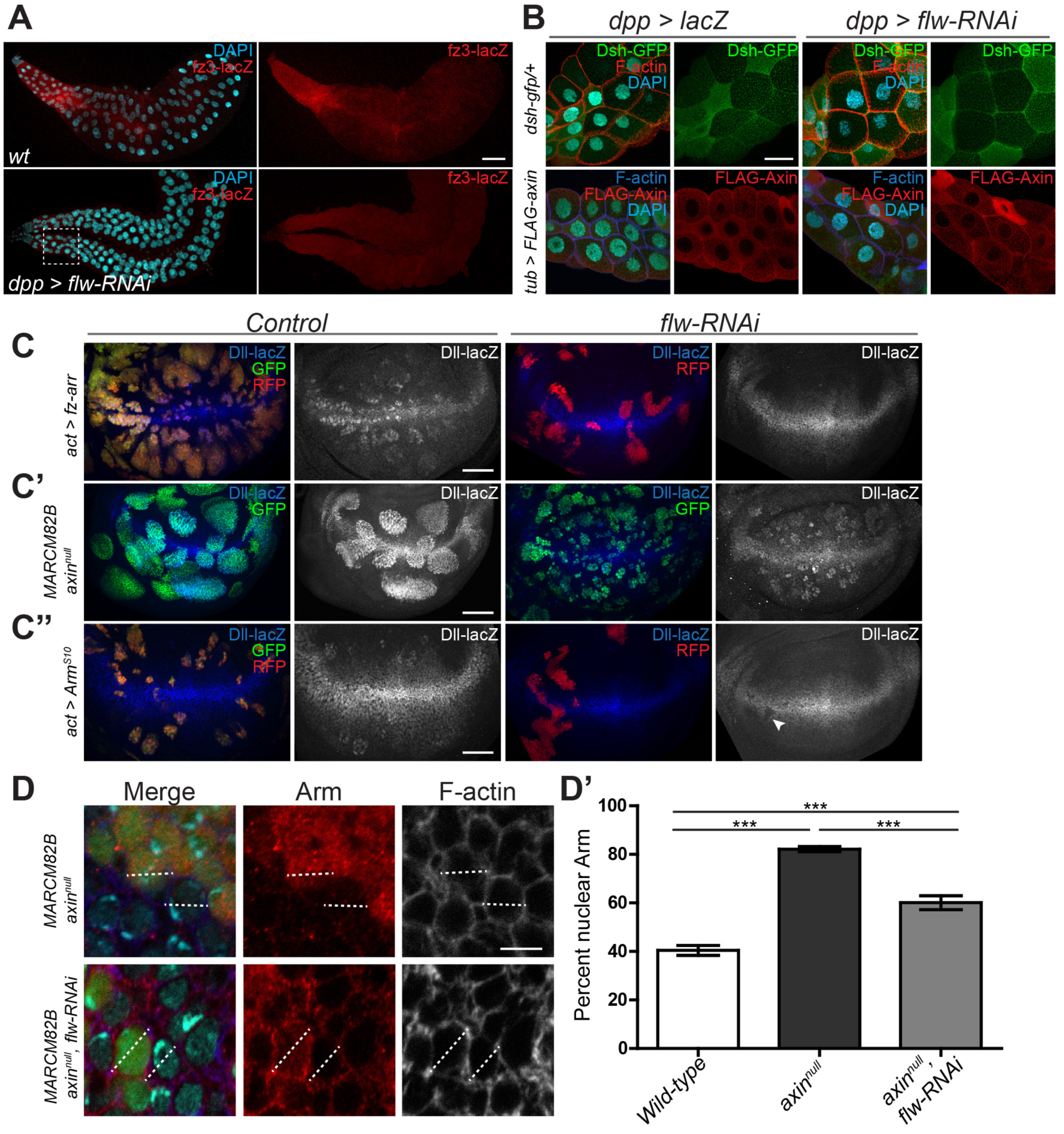
NMII activity inhibits Wg activation by reducing nuclear Arm independently of the destruction complex. (A) Salivary glands from control or *dpp>flw-RNAi* stained for *fz3* expression. (B) Localization of Dsh-GFP and FLAG-Axin in the proximal cells of the salivary gland, identified in the dashed line area of (A), (C-C”) Effects of *flw-RNAi* on ectopic *Dll-lacZ* in wing imaginal discs: (C) RFP marked flip-out clones expressing Fz-Arr and GFP *or flw-RNAi* (C) GFP-positive *axin^nul1^* MARCM clones *and flw-RNAi* in *axin^nul1^* MARCM clones. (C”) Arm^S10^ flip out clones with GFP *or flw-RNAi.* (D,D’) Effects of *flw-RNAi* on Arm distribution in GFP-marked *axin*^*null*^ cells. (D) DAPI was used to identify nuclei, and F-actin to mark the edges of the cell. (D’) Percent of nuclear Arm in cells was measured as an intensity plot (dotted line D) in wild type (n = 16), *axin*^*nul1*^ (n = 20), and *axin*^*null*^, *flw-RNAi* cells (n = 15). Data presented as mean ± SEM; *\*\*\*p<* 0.001. Scale bars: (A) 100 μm, (B-C”) 50 μm, (D) 5 μm.

To determine where NMII acts within the pathway, we induced ectopic Wg signaling at different points within the signaling cascade and asked if NMII could suppress ectopic target gene activation. An activated Fz-Arrow fusion protein induced ectopic *Dll* expression, which could be suppressed by *flw-RNAi* (Fig 2C), again suggesting that increased NMII inhibits Wg activity below the level of the receptors. To determine if NMII affects the destruction complex itself we generated *axin*^*null*^ MARCM clones, as Axin is the scaffolding protein on which the destruction complex assembles [27]. Clones lacking Axin had high levels of ectopic *Dll* and were large due to increased proliferation (Fig 2C). Expression o *flw-RNAi* in *axin*^*null*^ MARCM clones could not suppress ectopic *Dll*, but clones were generally smaller and did not show the smoothed edges seen in the *axin*^*null*^ clones (Fig 2C), suggesting that NMII can affect aspects of the *axin*^*null*^ phenotype. We next tested degradation resistant Arm^s10^[28]*. flw-RNAi* suppressed ectopic and even some endogenous *Dll* expression in clones with Arm^S10^ (Fig 2C”).

Cells with increased NMII activity had decreased levels of cytoplasmic and nuclear Arm (Fig 1H), but increased Arm at the apical surface (Fig 1H’), suggesting that NMII could suppress Arm^S10^ activity by inhibiting its ability to enter the nucleus or be retained there. To confirm this we looked at the relative distribution of Arm in *axin*^*null*^ tissue, to eliminate any variables NMII may have on destruction complex effectiveness and Arm turnover rates. Using F-actin to mark the edges of individual cells and DAPI to stain nuclei, intensity plots were drawn across individual cells to look at the distribution of Arm (Fig 2D dotted line). *axin*^*null*^ cells had roughly double the amount of Arm in the nucleus as wild type (Fig 2D’). Introduction of *flw-RNAi* resulted in a significant decrease in nuclear Arm in an *axin*^*null*^ background, but which was still higher than in wild type cells (Fig 2D’). These results are consistent with the level of Wg activity and maintained ectopic *Dll* seen in *axin^null^* cells with *flw-RNAi* (Fig 2C’). These findings suggest that increased NMII activity results in reduced entry or retention of Arm in the nucleus to initiate target gene transcription.

### NMII activation increases retention of adherens junction proteins

The regulation of Arm localization by NMII could be mediated by any one of the processes that NMII normally influences, including other downstream signaling pathways. We systematically examined key cellular functions of NMII to determine how NMII can modulate Wg signaling.

Loss of myosin phosphatase and increased NMII activity can stimulate JNK [*Drosophila* Basket (Bsk)] activity in the developing wing disc [29], and JNK has been shown to promote Wnt signaling (Wu et al., 2008). Using *dpp-Gal4* expressed along the anterior/posterior (A/P) boundary of the wing disc (Fig S4A,E) to express *flw-RNAi* reduced *Dll* expression, but in this context did not cause elevated JNK activity, seen by expression of JNK target gene *puc* [31](Fig S4A,A’,C,E,E’,G). Expression of a dominant negative Bsk^DN^, inhibiting JNK, did not affect *Dll* or *puc* in the wing pouch (Fig S4B,F). Importantly when co-expressed with *flw-RNAi*, *Dll* was still reduced and *puc* was unaltered (Fig S4D,H), indicating that NMII’s ability to suppress Wg signaling is not mediated through JNK. We next investigated NMII’s role in controlling integrin clustering for the formation of focal adhesion and ECM attachment (Vicente-Manzanares et al., 2009). Loss of function clones for the sole *Drosophila* β_PS_ integrin subunit [32] had no effect on expression of the Wg target Sens (Fig S4I).

Engl et al. (2014) studied the dynamics of NMII in suspension cell doublets. Following NMII activation, cells begin to constrict and pull away from one another, causing an influx and retention of E-cad to the AJ along with other proteins, including β-cat/Arm, to maintain and reinforce cell-cell adhesion. The increased apical Arm in cells with elevated NMII activity suggested that Wg signaling may be modulated by NMII’s ability to control E-cad clustering and retention during cell constriction. In clones expressing activated NMII (Sqh^EE^), constricting cells had increased *Drosophila* E-cad (DE-cad) and Arm along the apical surface at the AJ (Fig 3A, arrowheads). Activated myosin did not affect transcription of these genes, as seen by lacZ reporters (Fig 3B), suggesting that Arm and DE-cad were accumulating at the AJ in cells with increased NMII activity. Such an effect was previously shown, although no link to Wg signaling was tested [34,35]. We performed Fluorescence Recovery After Photobleaching (FRAP) analysis to measure the turnover of DE-cad and Arm at the AJ. FRAP of ubiquitously expressed DE-cad::GFP or Arm::GFP was measured along cell interfaces in wild type cells (Fig 3C,D, green box), cells expressing elevated NMII via *flw-RNAi* (Fig 3C,D, red box) and at the interface of wild-type*/flw-RNAi* cells (Fig 3C,D, blue box). FRAP revealed DE-cad recovery rates are significantly reduced at *any flw-RNAi* cell interfaces (Fig 3C) and contained a significantly higher immobile fraction (Fig 3C”), while Arm was unaltered (Fig 3D’,D”). The altered DE-cad recovery and immobile fraction rate, unaltered transcription yet protein accumulation is likely due to the increased force generated by NMII which can stabilizes cortical F-actin, which in turn stabilizes DE-cad at the AJ [33-36]. Accumulation of DE-cad at the AJ can subsequently bind and accumulate Arm.

**Fig 3.**
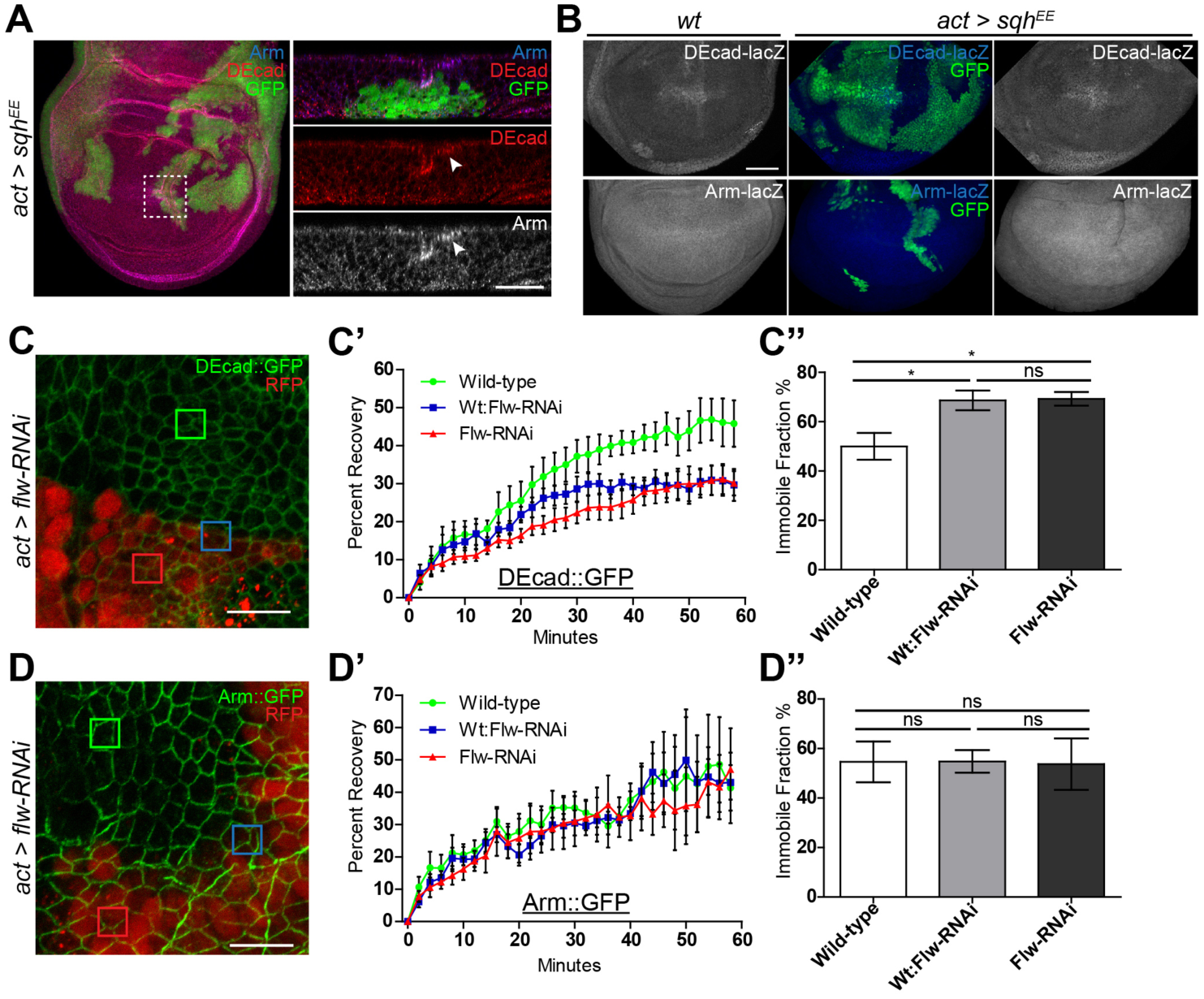
NMII activation increases retention of adherens junction proteins. (A-B) GFP-marked actin flip-out clones driving *sqh^EE^* stained for (A) DE-cad and Arm (arrowheads identify apical increases),(B) expression of *DE-cad* or *arm.* (C-D”) FRAP analysis of DE-cad::GFP and Arm::GFP in wing imaginal discs with RFP-marked *flw-RNAi* expressing flip-out clones. (C,D) DE-cad::GFP and Arm::GFP wing imaginal disc with squares indicating bleached regions of the wing disc. Green squares represents wild type cell interfaces, Blue for *wt.-flw-RNAi* cell interface, and Red for *flw-RNAi* cell interfaces. (C) AJ DE-cad::GFP (n = 6) recovery curves of FRAP analyses show cells adjacent to or expressing *flw-RNAi* have significantly slower recovery rates than wild type *(P* = 0.0029), (C”) and greater immobile protein fractions, *\*P<* 0.05. (D’) *flw-RNAi* had no effect on AJ Arm::GFP (n = 6) recovery curves from FRAP analyses *(P* = 0.4794), (D”) or immobile protein fractions. Data presented as mean and mean curve ± SEM. Scale bars: (A) 20 μm, (B) 50 μm, (C,D) 10 μm.

### NMII mediates DE-cad accumulation and sequesters Arm to the AJs, inhibiting Wg signaling

To further study the role of DE-cad in regulation of Arm, we expressed full length and a truncated version of DE-cad lacking the Arm binding domain (DEcadΔβ). Ectopic wild-type DE-cad was uniformly enriched along the cell periphery at the AJs, while DEcadΔβ expression was also seen in puncta (Fig 4A), since DE-cad unable to bind Arm is endocytosed and accumulates in vesicles [37]. Expressing either transgene had no effect on levels or distribution of F-actin (Fig 4B). However, ectopic DE-cad dramatically increased Arm at the AJ (Fig 4B), and could strongly suppress *Dll* expression (Fig 4C). DE-cad’s ability to suppress Wg signaling has been previously reported [38], and is likely due to the higher binding affinity of Arm to DE-cad over TCF binding for transcriptional activation [39]. Any cells exhibiting increased levels of DE-cad will titrate freely available Arm to the AJ, stabilizing DE-cad and increasing cellular adhesion. This is supported by the fact that expression of DEcadΔβ resulted in decreased levels of AJ Arm and did not suppress *Dll* expression (Fig 4B,C).

**Fig 4.**
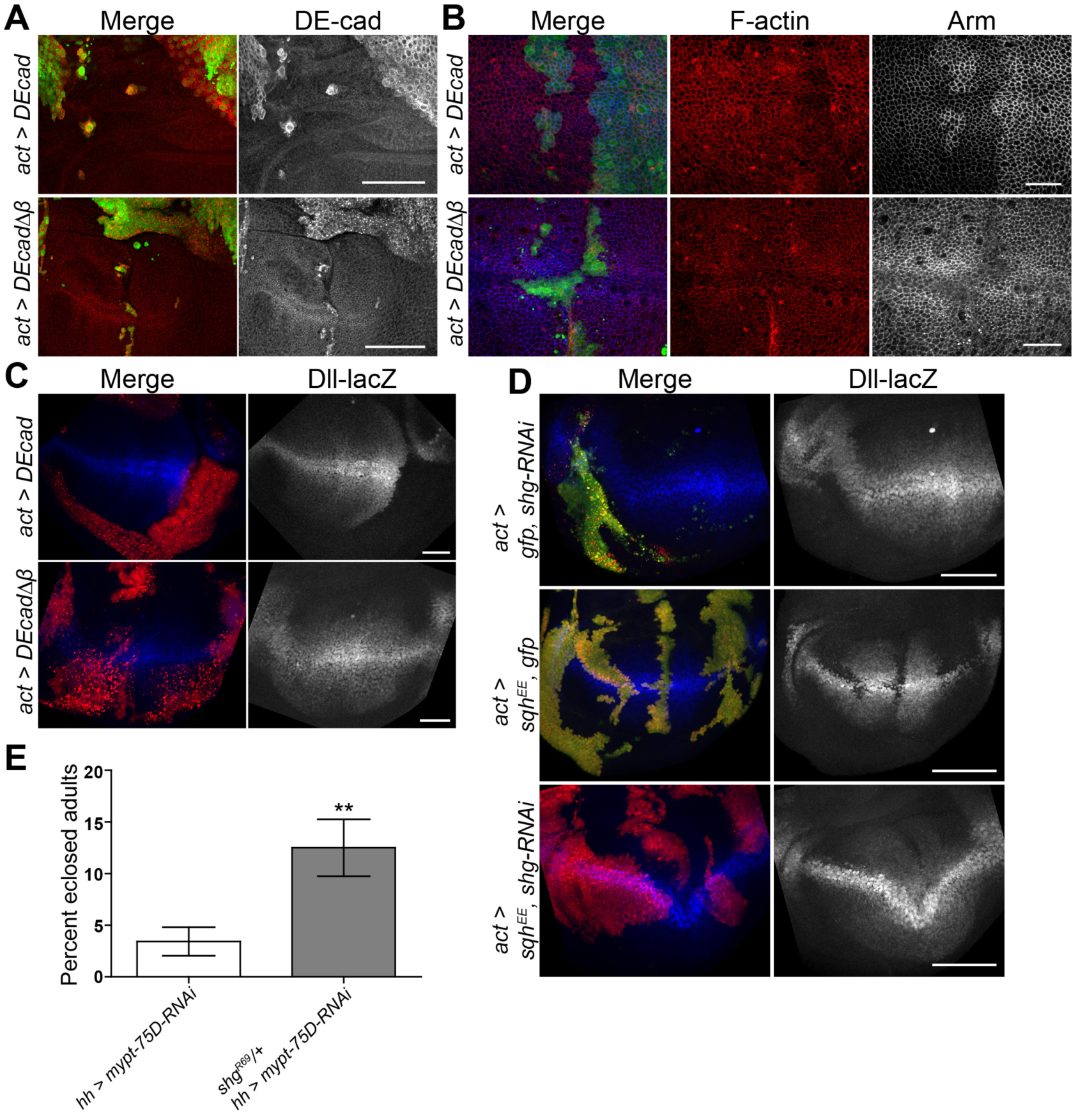
NMII activation inhibits Wg signaling through DE-cad. (A,B) GFP-marked actin flip-out clones expressing *DE-cad* or *DE-cad*Δ*β,* stained for (A) DE-cad, or (B) F-actin and Arm. (C) RFP-marked actin flip-out clones expressing *DE-cad* or *DE-cad*Δ*β,* stained for *Dll* expression. (D) *Dll-lacZ* expression in RFP-marked clones of the indicated genotypes. (E) Eclosion percentage of *hh>mypt-75D-RNAi* and *hh>mypt-75D-RNAi* heterozygous for *shg Drosophila.* Data presented as mean ± SEM; *\*\*P =* 0.0022; n ≥ 145. Scale bars: (A,C,D) 50 μm, (B) 20 μm.

We next examined the effects of reduction of overall DE-cad on Wg. RNAi against *shotgun (shg),* which encodes DE-cad, could induce cell death, but had no apparent effect on *Dll* (Fig 4D). Activated NMII (Sqh^EE^), caused a strong suppression of *Dll,* which was rescued by co-expression o *shg-RNAi* (Fig 4D), and *shg-RNAi* with Sqh^EE^ did not induce widespread cell death. Finally, *hh>mypt-75D-RNAi* adult flies had a much lower than expected viability and eclosion rate, which was rescued by heterozygosity for the *shg*^*R69*^ null allele [40](Fig 4E). These results suggest that elevated NMII can inhibit Wg activity through the accumulation of DE-cad, resulting in the titration of Arm to the AJ.

### NMII’s effect on the Wg pathway is mediated through F-actin stability

Engl et al. (2014) demonstrated that increased NMII activity results in decreased F-actin turnover, which can guide E-cad clustering and retention [34]. To confirm if F-actin levels in a developing tissue could also affect Wg activation we used the formin protein Diaphanous (Dia) which promotes the polymerization of filamentous actin [41]. Mitotic clones expressing a constitutively active Dia protein lacking its autoinhibitory domain (DiaΔDAD) had dramatic increases in levels of F-actin (Fig 5A-A”), and phenocopied the effects of increased NMII activation. Clones had decreased levels of *Dll* (Fig 5A, arrowhead), increased Arm and DE-cad along the apical surface of the cells (Fig 5A’ arrow, A”, open arrowhead), and cells in larger clones began to constrict, possibly due to filament cross-linking and bundling [42] (Fig 5A”, open arrowhead). To confirm if the effect of NMII on Wg is directly mediated through F-actin stability and constriction, we tested if reduced F-actin could alleviate NMII’s suppression of Wg signaling, using *dia-RNAi.* Cell with low levels of Dia had lower levels of F-actin, as well as increased apical cell surfaces, marked by Arm (Fig 5B). Reduction of F-actin via *dia-RNAi* had no major effect on endogenous Wg signaling, as seen by wild type *Dll* expression (Fig 5C). However, in an activated myosin background (Sqh^EE^), which strongly inhibits *Dll,* the co-expression of *dia-RNAi* could restore wild type *Dll* expression (Fig 5C). These results confirm that NMII can influence Wg pathway activity through its function to bind and stabilize actin leading to accumulation and constriction of filamentous actin.

**Fig 5.**
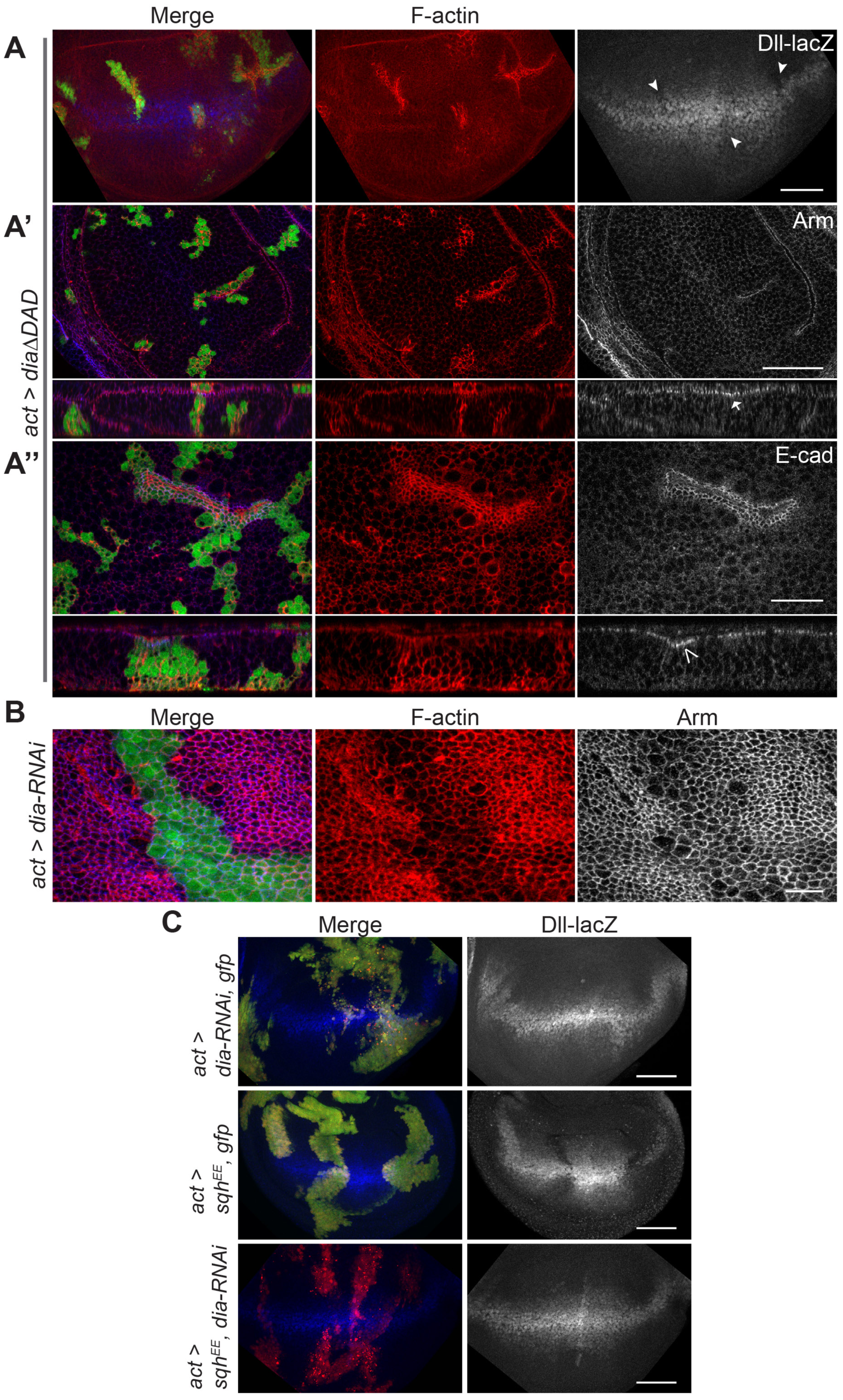
NMII inhibits Wg signaling by increased F-actin. (A-A”) GFP-marked actin flip-out clones expressing *dia*Δ*DAD* stained to detect (A) F-actin and *Dll-lacZ* (arrowheads indicate loss), (A’) F-actin and Arm and (A’’) F-actin and DE-cad. *diaΔDAD* induces increased apical AJ Arm (A’ arrow) and DE-cad (A” open arrowhead), and cell contractions (A” open arrow head). (B) GFP-marked actin flip-out clones expressing *dia-RNAi* stained for F-actin and Arm. (C) GFP-marked actin flip-out clones of indicated genotypes stained to detect *Dll-lacZ.* Scale bars: (A,A’,C) 50 μm, (A”) 20 μm, (B) 10 μm.

### NMII regulates Wnt in mammalian cells by sequestering β-cat to the AJs

We next examined the effects of increased NMII activation on Wnt signaling in human cell lines. Wnt pathway activity was induced in MCF7 and RKO cells by transfection of Wnt3A, and the response was measured using a TCF-responsive TOPFLASH transcriptional reporter [43]. MCF7 cells are epithelial, polarized and have well defined adherens junctions [44], while RKO cells are mutant for E-cad and completely lack adherens junctions (Fig 6C). The only β-cat present in RKO cells is solely for the regulation of Wnt activation [45]. Transfection of Wnt3A resulted in a significant increase in reporter activity in both cell lines, although RKO cells had a more robust response (Fig 6A,B). To increase NMII activation within these cell lines we transfected in siRNA against individual myosin phosphatase components. siPP1β could reduce total PP1β by ∽70% (Fig 6C), while siMYPT3 reduced MYPT3 (the ortholog of Mypt-75D) by ∽60% (Fig 6C). Knockdown of these components reduced the amount of the other myosin phosphatase protein, suggesting complex formation is essential for stability (Fig 6C). In MCF7 cells, knockdown of PP1β or MYPT3 could reduce Wnt activation significantly, and cotransfection of siPP1β and siMYPT3 reduced transcriptional activity back to baseline levels similar to cells with no Wnt3A (Fig 6A). In RKO cells there was no significant change, in fact there was a slight increase in Wnt transcriptional activation (Fig 6B). These results indicate that increased NMII activation from reduction of myosin phosphatase components can inhibit Wnt activity in mammalian cells as well, but only in cells that contain adherens junctions.

**Fig 6.**
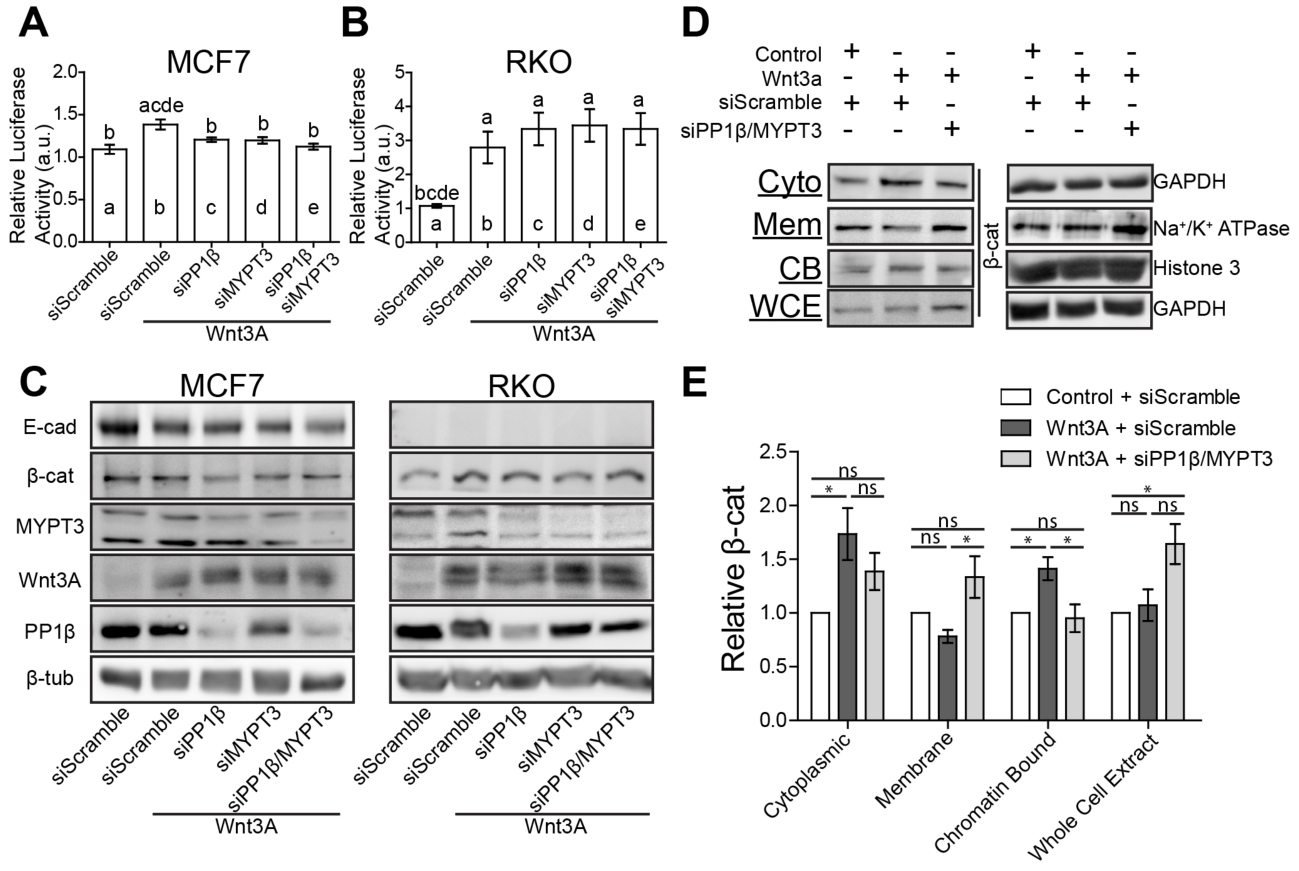
NMII activation recruits β-cat to cell membranes inhibiting Wnt signaling. (A,B) Wnt-responsive luciferase TOPFLASH assay measuring TCF/LEF reporter activity in (A) MCF7 and (B) RKO cells following Wnt3A transfection and siPP1β and siMYPT3 transfection. Data are presented as mean ± SEM with letters above representing significant difference from corresponding column, *(P<0.01).* (C) Western blot analysis of total cellular levels of E-cad, β-cat, MYPT3, Wnt3A and PP1β. β-tub was used as a loading control. (D) Western blot analysis of β-cat in cytoplasmic (Cyto), membranous (Mem), chromatin-bound (CB), and whole cell extract (WCE) fractions in MCF7 cells. GAPDH, Na^+^/K^+^ ATPase, and Histone 3 were used as loading controls for corresponding fractions. Complete fraction, see Fig S5. (E) Quantification of relative levels of β-cat from each fraction taken from (D), normalized to Control + siScramble conditions. Data shown as mean ± SEM (n = 4 biological replicates), ns = not significant, *\*P<0.05.*

Although reduction of PP1β or MYPT3 was able to inhibit Wnt signaling in MCF7 cells there was no dramatic change in overall β-cat levels (Fig 6C), suggesting that there may be a localization defect as seen in *Drosophila.* Transfection of Wnt3A induced a significant increase of cytoplasmic and chromatin-bound (transcriptionally active) β-cat, while membrane associated (AJ) β-cat slightly decreased (Fig 6D,E). The decrease in membranous β-cat is likely due to the fact that Wnt can induce mild EMT effects in MCF7 and other epithelial cancer lines [46]. Similar effects were seen with E-cad (Fig S6). Whole cell extracts only showed a minor increase in total β-cat (Fig 6D,E). A striking inverse in distribution was seen after increasing NMII activation via siPP1β and siMYPT3 transfection. Cytoplasmic levels of β-cat decreased, and there was a significant reduction in chromatin-bound levels back to baseline, matching results seen in TOPFLASH assays (Fig 6A,D,E). Cells with activated NMII had a significant increase in membrane associated β-cat levels, as well as increased total β-cat (Fig 6D,E). Considering the increase in total β-cat, yet lack of transcriptional activation, and redistribution of the protein within the cell, we propose that in mammalian cells that have elevated NMII activity, titrate freely available β-cat to the adherens junctions to enforce cell-cell adhesion at the cost of transcriptional activation of Wnt targets.

### NMII activation modulates Wnt signaling during development and homeostasis to maintain cell-cell adhesion

We next tested this model in *Drosophila.* We generated mitotic recombinant clones that can form and maintain AJs without the need for Arm using functionally validated fusion proteins in which DE-cad is fused to α-cat (DEcad::αCat), as well as a truncated fusion of the proteins lacking their Arm binding domains (DEcadΔβ::αCatΔVH1) [47]. Actin flip-out clones expressing either transgene led to increased levels of DE-cad and did not affect levels of F-actin (Fig S6A-D). DEcad::αCat could still bind Arm, forming puncta within the cells, and resulting in a suppression of *Dll* expression (Fig S6A,C). DEcadΔβ::αCatΔVH1 did not affect Arm distribution or *Dll* expression (Fig S6B,D), so we utilized this transgene for further experiments.

Mitotic clones of the *shg*^*R69*^ null allele are non-viable and were quickly extruded from wing disc (data not shown). When DEcadΔβ::αCatΔVH1 was expressed in these cells, clones could divide and grow (Fig 7A), confirming that DEcadΔβ::αCatΔVH1 could form AJs without binding Arm. Clonal tissue had normal Dll protein levels and F-actin (Fig 7A). The expression of *mypt-75D-RNAi* in *shg* clones had no effect on Dll, but did induce minor accumulations in F-actin (Fig 7A’, arrowheads). These results were mimicked when DEcadΔβ::αCatΔVH1 was co-expressed with Sqh^EE^, namely clonal tissue still constricted and had elevated F-actin, but the reduced *Dll* expression was significantly rescued (Fig S6E, Fig 4D). These results indicate that NMII activation can only suppress Wg signaling in this epithelium when DE-cad binds to Arm.

**Fig 7.**
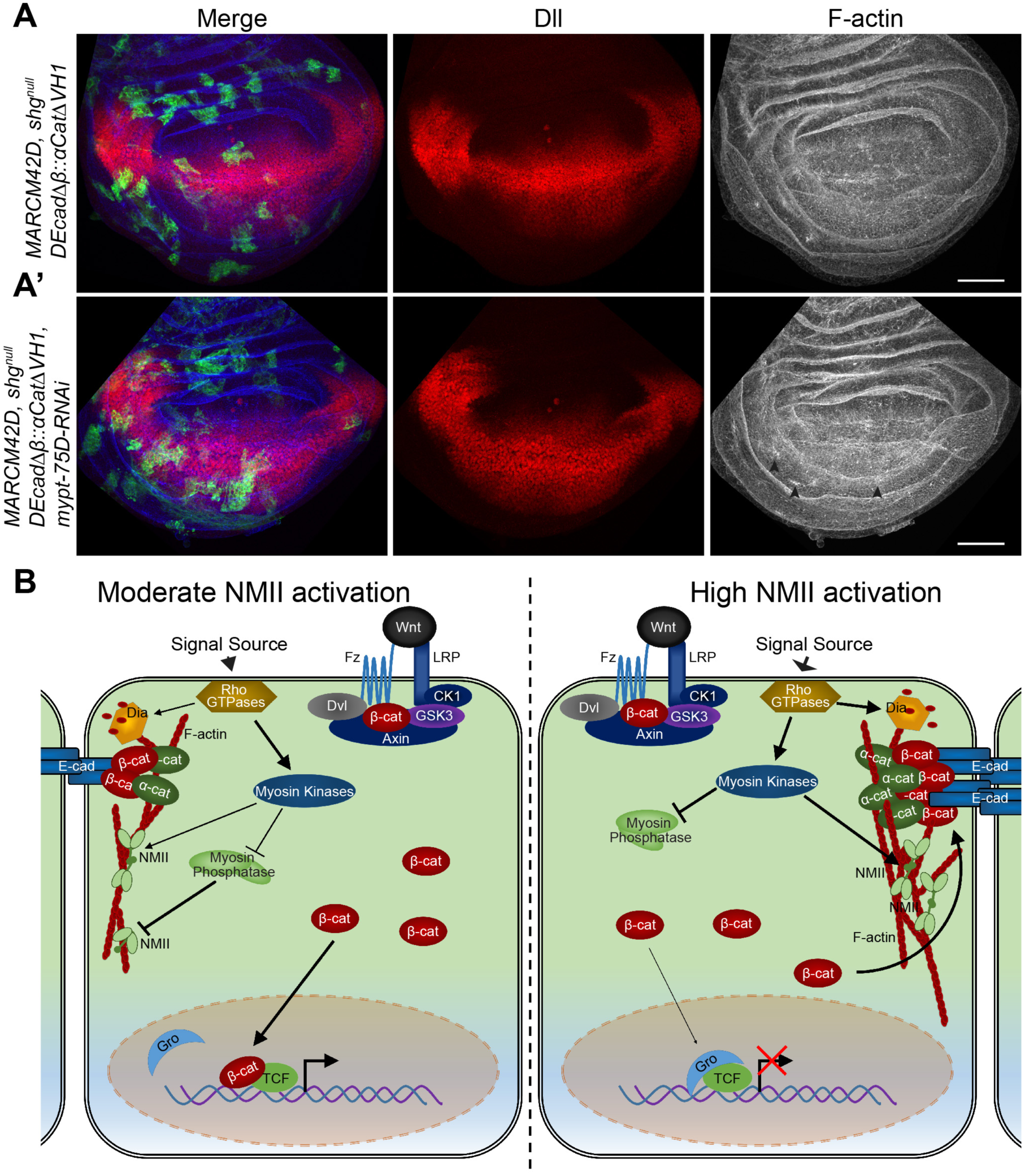
NMII activation inhibits Wnt signaling in a dynamic fashion across developing tissue by Arm titration to AJs. (A,A) GFP-marked *shg*^*null*^ MARCM clones expressing *(A) DK-caaΔβ::a-catΔVH1* and (A) *DE-cad*Δ*β::α-catΔVH1* with *mypt-75D-RNAi,* stained for Dll and F-actin (arrowheads show F-actin accumulation). (B) Model for regulation of Wnt signaling by activation of NMII resulting in AJ accumulation and stabilization in response to contractile forces; see text for details. Scale bars: (A,A’) 50 μm.

## Discussion

The link between mechanical forces and canonical Wnt signaling has been extensively studied in contexts such as mesoderm differentiation during cardiomyogenesis [48] and in tissue stiffness of the ECM in stem cell behavior or carcinogenesis [7,9]. Less research has focused on how mechanical forces may directly influence Wnt activation in normal developing epithelia. Our work shows that increased NMII activation in epithelial cells induces contraction and accumulation of cortical F-actin, and as a result E-cad accumulates and titrates freely available Arm/β-cat to the AJs in order to maintain cell-cell adhesion. The resulting decreased levels of cytoplasmic Arm/β-cat causes insufficient nuclear translocation and reduced Wnt target gene transcription (Fig 7B).

We show that NMII activation can inhibit the nuclear accumulation of Arm causing a suppression of overall transcriptional initiation, even in genotypes lacking Arm degradation machinery or expressing Arm resistant to degradation. These results identify a distinct mechanism in regulating Wnt signaling independent of the adhesion-based enhanced destruction complex activity turnover of Arm/β-cat [49], and were complementary to those of Greer et al. (2013), who identified that RhoGEF and GTPase activity (upstream activators of NMII) could suppress Arm localization or activation in developing *Drosophila.* We further excluded the possibility that NMII suppresses Wnt through several known interaction mechanisms, like JNK and ECM/Integrin signaling [9,50].

Cells with elevated NMII activity and nearby adjacent cells that were under increased contraction exhibited elevated Arm and DE-cad at AJ, while transcription rates of the genes encoding these proteins appeared normal. DE-cad (*shg*) was previously identified as a Wnt target gene in wing discs [21]. Its maintained transcription in *flw-RNAi* cells may be due to compensation by another transcriptional mechanism. Furthermore, the recovery rate of DE-cad to the AJ was reduced, and contained a higher immobile fraction of DE-cad, indicating accumulation and retention. Although we did not detect any significant changes in Arm recovery rates or immobile fraction at the AJs, this may be due to positional effect of our measurements across the wing disc. Total Arm, and its distribution within a cell vary dramatically across the wing disc. In order to maintain healthy tissue to generate accurate measurements, our *flw-RNAi* clonal induction at random positions may explain the high variance. As FRAP experiments encompassed the entire AJ and membrane, the recovery rates are likely an indirect measure of vesicle trafficking of Arm and DE-cad [36]. This is bolstered by the fact that decreased actomyosin levels have been shown to increase AJ endocytosis in developing *Drosophila* [36]. The accumulation of E-cad is likely an active mechano-sensitive mechanism to bolster cell-cell adhesion, potentially through plasma membrane clustering and vesicle-based redistribution [5,33,34].

Early studies of *arm* in Drosophila revealed the dual role of β-catenin in both Wg signaling and at the AJ through interactions with E-cad and α-catenin [51]. Elevated E-cad has been previously shown to suppress Wg signaling [38], but no defined mechanism was identified. We propose, like others before, that this effect is likely due to the higher binding affinity of Arm/β-cat to E-cad over TCF for transcriptional activation [39,52]. In this study we reveal that the activation status of NMII can directly impact the stabilization of the β-cat/E-cad complex and thus affect Wnt signaling readouts. We validated this model by expressing wild type DE-cad or mutant DE-cad, lacking the Arm binding sequences. Our results demonstrated that ectopic DE-cad caused high levels of Arm to be enriched along the AJ, and strongly suppressed Wg target gene expression, while mutant DE-cad led to reduced AJ Arm and did not affect Wg targets. Importantly neither of these transgenes had any effect on F-actin, showing that E-cad acts downstream of F-actin in this NMII activation pathway to suppress Wnt signaling. In addition the reduction of DE-cad in wing disc tissue was able to rescue Wg activity defects and viability of flies expressing activated NMII.

In our *in vivo* work, we were able to confirm the model by Engl et al. (2014) and Hong et al. (2013) that NMII activation recruits and stabilizes DE-cad, Arm, and other AJ core proteins in developing tissue by stabilization and accumulation of F-actin, and in our context resulting in a suppression of Wg activation. The expression of a constitutively active Formin protein phenocopied the ability of activated NMII to inhibit Wg target gene expression, and the reduction of F-actin was able rescue the effect of increased NMII activity on Wg target gene expression.

Building on our *Drosophila* work, we were able to confirm that NMII has similar effects on Wnt in human cells that contain AJs. Stimulation of NMII was able to suppress Wnt transcriptional activation in polarized MCF7 cells containing AJ, but had no effect in RKO cells lacking E-cad. Importantly, increased NMII activity following knockdown of myosin phosphatase induced a significant redistribution of β-cat out of the chromatin-bound fraction to the membrane, and increased overall β-cat protein levels within the cells. Although these cells had significantly higher levels of β-cat, their inability to initiate Wnt target gene transcription indicates that β-cat is being sequestered to the AJ. We validated this model by generating clonal tissue in which we replaced endogenous E-cad with a fusion protein that does not require Arm/β-cat for the formation of complete AJ. In these cells, when NMII activity was stimulated there was no suppression of Wnt target gene transcription, confirming NMII inhibits Wnt by titrating Arm/β-cat to the adherens junctions.

This may be a physiological regulatory mechanism in developing tissue for proliferation, patterning, and morphogenesis, and later in the homeostasis of epithelia. As tissues proliferate, change shape, and respond to physical cues, the cells respond to all these factors and induce variable levels of NMII activation. In order to maintain overall tissue integrity as cells change their shape, cell-cell adhesion must be increased. This results in the sequestration of Arm/β-cat to increase AJ adhesion and inhibit canonical Wnt’s ability to promote patterning. In essence the preservation of tissue integrity overrides Wnt-inducible gene expression in epithelia. Recently there have been comparable instances of this in other developmental signaling pathways, through distinct mechanisms. Increased NMII activation and cytoskeleton tension have been widely identified to inhibit Hippo signaling, while stimulating JNK activation in epithelia [29,53,54]. With this study, we demonstrate that canonical Wnt signaling is another key developmental pathway that is regulated by NMII.

It will be interesting to determine if these dynamic contractile forces exerted upon developing tissue are also critical for the maintenance of stem cell niches. For instance in intestinal crypts, Wnt activity is refined to the very basal cells of the crypt for the maintenance of stem cells, where cells are apically constricted to form the concave base of the crypt [55]. Monitoring Wnt activity in crypt cells as well as proliferation and differentiation rates when exposed to increased or decreased levels of NMII activity, or even removing the tissue to grow on a flat surface, may provide insights into the relevance of why and how tissue structures arise due to the forces that are exerted on cells across a tissue in order to regulate homeostasis. The modelling of stem cell niche formation in crypts has suggested that the loss of curvature regulation can result in tissue consisting of Paneth cells of the crypt and undifferentiated cells, which is seen in some cases of intestinal adenoma and carcinoma [55]. Our results here have provided new and supportive evidence that the interactions between biochemical signaling, Wnt in this case, and mechanical forces, guiding cell shape and adhesion, are critical for the normal development and maintenance of healthy epithelial tissue in an organism.

## Materials and Methods

### Drosophila husbandry, crosses, and clone generation

Fly strains and crosses were raised on standard medium at 25°C unless stated otherwise. *w^1118^* was used as wild type. In assays examining the interactions between two or more UAS transgenes, control crosses were performed with *UAS-lacZ* or *UAS-GFP*, to rule out effects due to titration of Gal4. Heat-shock inducible actin flip-out clones were generated by crossing either RFP-marked flip-out or GFP-marked flip-out strains to corresponding lines, then larvae were heat-shocked at 37°C for 12.5 or 15 minutes respectively, 48-72 hours after egg laying (AEL) (depending on the assay), and incubated at 29°C until dissection. Mosaic analysis with a repressible cell marker (MARCM) clones were generated by crossing MARCM lines to corresponding lines and larvae were heat-shocked at 37°C for 1.5 hours, 48 (*MARCM82B*) or 72 (*MARCM19A* and *MARCM42D*) hours AEL, and incubated at 29°C until dissection.

The following fly strains were used: (1) *Dll-lacZ* (BL10981), (2) *dpp-Gal4* (BL 1553),(3) *y*^1^ *sc* v*^1^; *P{TRiP.HMS00521}attP2* (*mbs-RNAi*) (BL 32516), (4) *UAS-sqn-RMAi* (BL 31542, 38222, 32439, 33892), (5) *UAS-bsk*^DN^(BL 6409), (6) *y*^1^ *w*^67c23^, *P {lacW}shg^k03401^/CyO (shg-lacZ)* (BL 10377), (7) *w*\*, *P{FRT*(*w*^*hs*^*)}G13 shg*^1^/*CyO*; *P{Ubi-p*63*E-shg.GFP}* (*ubi-shg-GFP*) (BL 58471), (8) *UAS-arm*^s10^(BL 4782), (9) *arm-GtP* (BL 8555), (10) *UAS-aiaΔDAD* (BL 56752), (11) *y*^1^ *v*^1^, *P{TRiP.HM05027}attP2* (*UAS-aia-RMAi*) (*BL* 28541), (12) *mys*^1^ *FRT19A/FM7c* (*β*_*PS*_^*null*^) (BL 23862), (13) *UAS-p35* (BL5072), (14) *w* sqnr^AX3^ P{neoFRT}*19*A*/*FM7c* (BL 25712) (obtained from the Bloomington Drosophila Stock Center), (15) *UAS-flw-RNAi)* (VDRC 104677, 29622), (16), *UAS-mypt-75D-RNAi* (VDRC 109909), (17) *UAS-mbs-RNAi* (VDRC 105762), (18) *UAS-shg-RNAi* (VDRC 27082), (obtained from the Vienna Drosophila Resource Center), (19) ;;*puc*^*E69*^-*lacZ* (Ring & Martinez Arias, 1993), (20) *fz3-lacZ*/*FM7a* (Sato et al., 1999), (21), *yw, arm-lacZ, FRT19A;; eyFLP/TM6B (arm-lacZ)* (Vincent et al., 1994), (22) *hh-GAL4* (Port et al.,2011), (23) *UAS-sqn*^E20E21^ *(UAS-sqn*^*EE*^) (Winter et al., 2001), (24) UAS-DEcad::αcatΔVH1 (Desai et al., 2013), (25) *shg*^*R*69^, *FRT42D*/*CyO* (*shg*^*null*^) (Godt & Tepass, 1998), (26) *hsFLP*;; *Act>CD2>Gal4, UAS-GFP/SM6∽TM6* (GFP-marked flip-out) (Bruce Edgar, Zentrum für Molekulare Biologie der Universität Heidelberg, Germany), (27)*hsflp*^122^, *tub-gal80*, *FRT19A*; *Act-Gal*4, *UAS-GFP;* (MARCM19A) (Rongwen Xi, NIBS, Beijing, China), (28*) y*, *w*, *hsflp*, *UAS*-*GFP*, *tub*-*Gal4*;;*FRT82B tubGal80* (MARCM82B) (Bruce Edgar, Zentrum für Molekulare Biologie der Universität Heidelberg, Germany), (29) *;;UAS-GFP,hsflp122,FRT42,tub-GAL80,tub-GAL4/TM6B* (MARCM42D) (Jessica Treisman, NYU School of Medicine, USA), (30) *UAS-DEcadΔβ::αcatΔVHJ* (Ulrich Tepass, University of Toronto, Department of Cell & Systems Biology), (31*) Dsh-GFP/CyO* (Jeffrey Axelrod, Dept. of Pathology, Stanford University School of Medicine, USA), (32) *yw; tub>FLAG-axin/CyO* (Marcel Wehrli, Oregon Health & Science University, USA), (33) *UAS-fz-myc-arr,* (Marcel Wehrli, Oregon Health & Science University, USA), (34) *axin*^*S044230*^, *FR182B*/*TM6B* (axin^*null*^) (Marcel Wehrli, Oregon Health & Science University, USA), (35) *en-Gal4,UAS-gfp* (Konrad Basler, Institute of Molecular Life Science, University of Zurich, Switzerland), (36) *y,w,hsflp122; sp/CyO; Act>CD2>GAL4,UAS-Rbr/lMoB* (RFP-marked flip-out), (37) *shg*^R69^, *FRT42D*,*mypt*-*75D*-*RNAi*; *UAS-DEcadΔβ::αcatΔVHl/SM6a∽TM6B*.

### Immunofluorescence, wing mounting and imaging

Third-instar larvae were dissected in phosphate-buffered saline (PBS). Wing imaginal discs and salivary glands were fixed in 4% paraformaldehyde at room temperature for 20 min followed by three washes in PBS for 5 minutes. Tissue was blocked [2% BSA diluted in PBS 0.1% Triton X-100 (PBST)] for 45 min at room temperature, followed by incubation with primary antibodies overnight at 4°C. Tissue was then washed three times for 5 minutes with PBST and incubated with secondary antibodies at room temperature for 1.5 hours. Phalloidin-rhodamine, or −647 (ThermoFisher Scientific), or Phalloidin-Fluorescein (Sigma-Aldrich) and DAPI (ThermoFisher Scientific) were added at this point, if required. A final series of three PBST washes were performed, followed by mounting in 70% glycerol in PBS. The following primary antibodies and dilutions were used: mouse anti-β-galactosidase (1:2000 Promega), mouse anti-Wg (1:100 DSHB), mouse anti-Arm (1:50 DSHB), rabbit anti-cleaved Caspase 3 (1:100 Cell Signaling), guinea pig anti-Sens (1:500, a gift from Hugo Bellen, Dept. of Molecular and Human Genetics, Baylor College of Medicine, USA), mouse anti-Dll (1:300, a gift from Ian Duncan, Dept. of Biology, Washington University in St. Louis, USA), rabbit anti-Phospho-Myosin Light Chain 2 (Ser19) (p-MyoII) (1:25 Cell Signaling), mouse anti-GFP (1:500, Cell Signaling), rabbit anti-FLAG (1:200, Sigma), rat anti-DEcad (extracellular domain) (1:50 DSHB), rat anti-Ci (1:50, DSHB). Secondary antibodies (Jackson ImmunoResearch) were used at a 1:200 dilution.

A minimum of 20 discs or glands were mounted per slide for a given genotype. Adult wings were dissected in 95% ethanol and mounted in Aquatex (EMD Chemicals). A minimum of 12 wings were mounted per genotype for analysis. Microscopy images were taken with an A1R laser scanning confocal microscope (Nikon) and adult wings were imaged with a Zeiss Axioplan 2 microscope.

### Live imaging and FRAP

Third-instar wing imaginal discs were dissected and mounted in SFX-Insect serum-free insect cell culture medium (Hyclone) supplemented with methyl cellulose (Sigma-Aldrich) at a concentration of 4% wt/vol to increase viscosity to prevent disc drifting while imaging. FRAP assays were carried out on an A1R laser scanning confocal microscope (Nikon), with 60× objective, 5× artificial zoom. GFP within the region of interest (ROI) was photobleached with a 405 nm and 488 nm UV laser at 100% power for 15 seconds. GFP recovery specifically along the AJs was then recorded by time-lapse imaging over 60 minutes at 2-minute intervals. Focal planes were maintained by manual focus during time-lapse. Any samples that exhibited phototoxicity or additional photobleaching in control regions during the time-lapse were excluded. GFP recovery rates of individual ROI data were normalized with pre-FRAP equal to 100% and post-FRAP equal to 0%. AJ immobile fractions of proteins were calculated as pre-FRAP fluorescence intensities, minus the end value of recovered fluorescence intensity of individual ROI.

### Image processing, measurements, and statistical analysis

Following image acquisition, images were processed using NIS Elements (Nikon) and Adobe Photoshop CS6. Immunofluorescence images are presented as maximum intensity projections of Z-steps spanning the entire tissue.

Distribution of Arm was determined by single line fluorescence intensity plot across individual cells with NIS Elements (Nikon). Cell edges were determined by peak F-actin fluorescence, and increased DAPI across the plot line marked the nucleus. Percentage of nuclear Arm, was determined as the value of Arm within the nuclear area over the total Arm across the intensity plot of the cell. Cell surface area was quantified using ImageJ (NIH) software. All data quantifications were performed in Microsoft Excel or GraphPad Prism and figures were made using Adobe Illustrator CS6.

Statistical analyses were performed using GraphPad Prism. Significant differences between two genotypes was determined by two-tailed Student’s t tests. One-way ANOVA was performed for multiple comparisons, with Tukey’s multiple comparison as a post-test. All quantified data are presented as mean ± SEM, and *P*< 0.05 was considered statistically significant. Significance depicted as * = *P*< 0.05, ** = *P*< 0.01, *** = *P*< 0.001, ns = not significant.

### Plasmid constructs and oligonucleotides

The following plasmids were used in this study: pCMV-Myc (control vector) (Clontech), TOPFLASH [43], FOPFLASH [43], pRL-CMV (Renilla Luciferase) (Promega), pcDNA-Wnt3A a gift from Cara Gottardi. The following siRNA oligonucleotides were used in this study: PPP1R16A (MYPT3): ID s39809 (ThermoFisher Scientific), PPP1CB (PP1β): ID s10935 (ThermoFisher Scientific), siRNA Negative control #1 (Scramble) (ThermoFisher Scientific).

### Cell culture

Cells were cultured in six-well plates at 37°C in 5% CO_2_. RKO (CRL-2577; ATCC) cells were grown in Dulbecco’s modified Eagle’s medium (DMEM; Gibco) supplemented with 10% heat-inactivated fetal bovine serum (hi-FBS; Invitrogen). MCF7 (HTB-22; ATCC) cells were grown in DMEM:F-12 (Gibco) supplemented with 10% hi-FBS (Invitrogen). Reverse transfections of siRNA complexes was performed in cells seeded at 50% confluence with Lipofectamine RNAiMAX (ThermoFisher Scientific) in Opti-MEM (Gibco), according to manufacturer’s instructions. 24 hours after seeding, transfections of plasmid DNA was performed with Lipofectamine 3000 and P3000 reagent (ThermoFisher Scientific), according to manufacturer’s instructions. When required, the final amount of DNA used for transfection was kept constant by the addition of control vector DNA. All cells were harvested 48 hours after transient DNA plasmid transfection for subsequent assays.

### Lysate collection and immunoblotting

Lysates of whole cell extract of MCF7 and RKO cells transfected with respective plasmids and siRNA were generated by collecting and treating the cells with cell lysis buffer (Cell Signaling Technology) supplemented with protease inhibitors (Roche). Lysates were then sonicated for several seconds on ice, followed by a 16,300 g centrifugation for 10 min at 4°C. The supernatant was removed and protein concentrations were determined by bicinchoninic acid (BCA) assay (ThermoFisher Scientific). MCF7 cellular fraction lysates were generated by the Subcellular Protein Fractionation assay (ThermoFisher Scientific) according to manufacturer’s instructions. Protein concentrations were equalized within individual fractions, as determined by BCA assay (ThermoFisher Scientific). Lysates were boiled for 10 min with Laemmli buffer, then separated on 8-12% SDS-PAGE gels. Proteins were transferred onto nitrocellulose membranes, and probed against the following primary antibodies: rabbit anti-β-catenin (1:1000, Cell Signaling), rabbit anti-E-cadherin (1:1000, Cell Signaling), mouse anti-PPP1R16A (MYPT-3) (1:500, abcam), mouse anti-Protein Phosphatase 1 beta (PP1β) (1:1000, abcam), rabbit anti-Wnt3A (1:1000, Cell Signaling), mouse anti-β-tubulin (1:1000, ABM), rabbit anti-GAPDH (1:3000, Cell Signaling), mouse anti-Na+/K+ ATPase (1:50 DSHB), rabbit anti-Histone H3 (1:1000 Cell Signaling). Membranes were visualized using the enhanced chemiluminescence (ECL) western blotting substrate (Pierce) with a LAS4000 luminescent imager (Fujifilm). The protein levels were determined using ImageJ (NIH) software to perform densitometry. Transfections and western blotting was performed in triplicate.

### Transcriptional reporter assay

Luciferase assays were performed in MCF7 and RKO cells with the Dual-Luciferase Reporter Assay System (Promega) according to manufacturer’s instructions. TOPFLASH or negative control FOPFLASH reporter gene plasmids, with the control reporter plasmid encoding Renilla luciferase (to normalize transfection efficiencies and for monitoring cell viability) were transfected with each expression vector as indicated, to determine overall Wnt pathway activity through TCF/LEF reporter activity. The values shown represent the mean ± SEM from four biological replicate transfections, performed each time in triplicate. TOPFLASH values were normalized to the FOPFLASH reporter activity equal to 1 for each individual transfections series.

## Acknowledgements

We thank the following individuals and stock centers for reagents, comments, and fly strains: Ulrich Tepass, Cara Gottardi, Ken Irvine, the Bloomington Drosophila Stock Center, the Vienna Drosophila RNAi Center and the Developmental Studies Hybridoma Bank.

## Supporting Information captions

**Figure S1.**
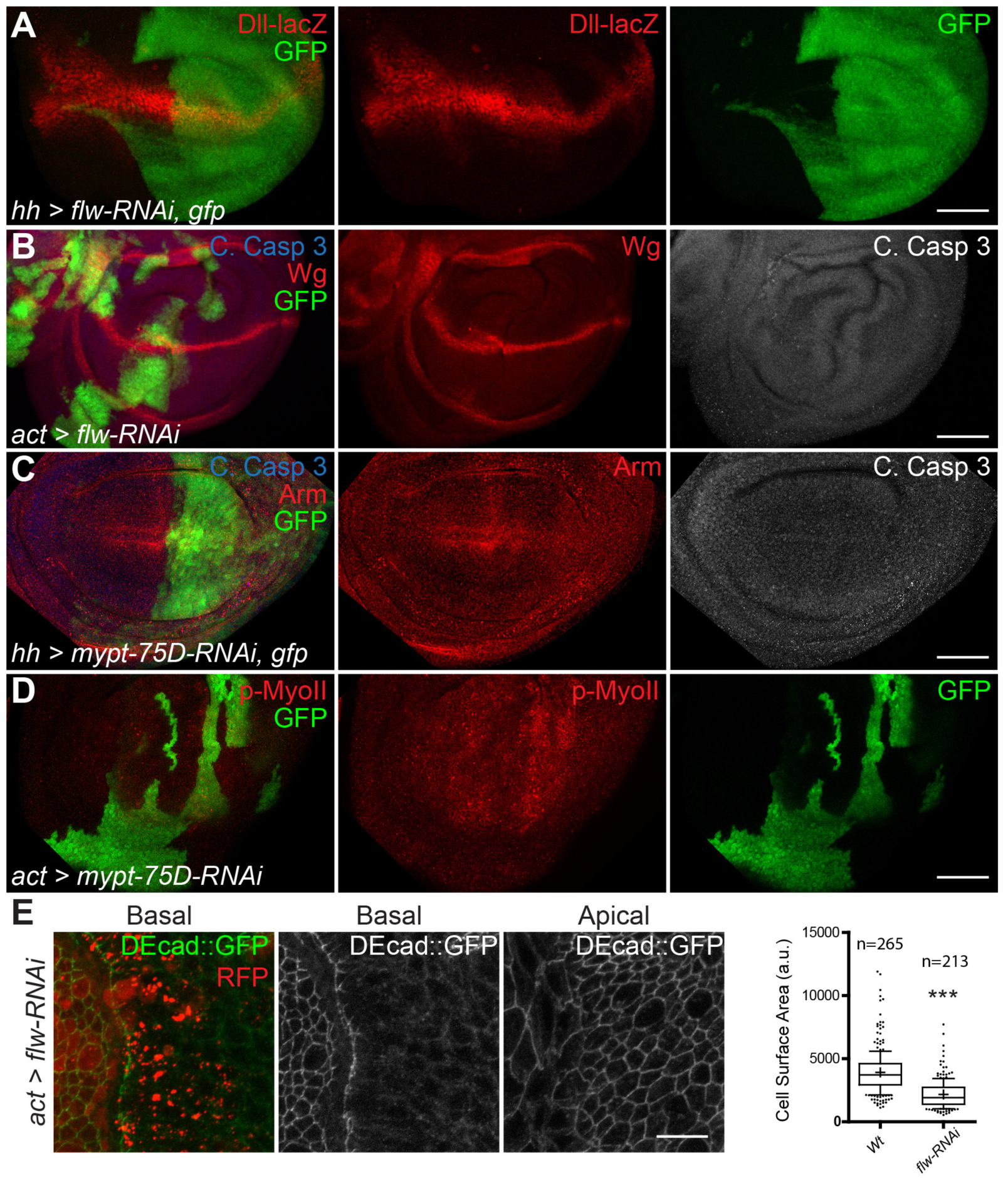
Knockdown of myosin phosphatase inhibits Wg activity, and promotes NMII activity during wing development. Related to Figure 1. (A) Wing discs in which *hh-Gal4* drives GFP and *flw-RNAi* in the posterior compartment, stained for *Dll-lacZ.* (B) GFP-marked actin flip-out clones *expressing flw-RNAi,* stained for Cleaved Caspase 3 (C. Casp3) and Wg. (C) Arm and C. Casp3 staining in discs with *hh-Gal4* driven *mypt-75D-RNAi* and *GFP.* (D) GFP-marked actin flip-out clones expressing *mypt-75D-RNAi,* stained for phospho-Myosin II (p-MyoII). (E) Cell surface areas of apical and basal sections of cells ubiquitously expressing DEcad::GFP fusion proteins to mark cell boundaries, with RFP-marked actin flip-out clones expressing*flw-RNAi*. Cell surface area data represented as box plots 25-75 percentile, whiskers 10-90 percentile, (-) median, (+) mean and (•) outliers. Wild-type (n=265) and *flw-RNAi (n=213)* cells, *** = *P*< 0.001.

**Figure S2.**
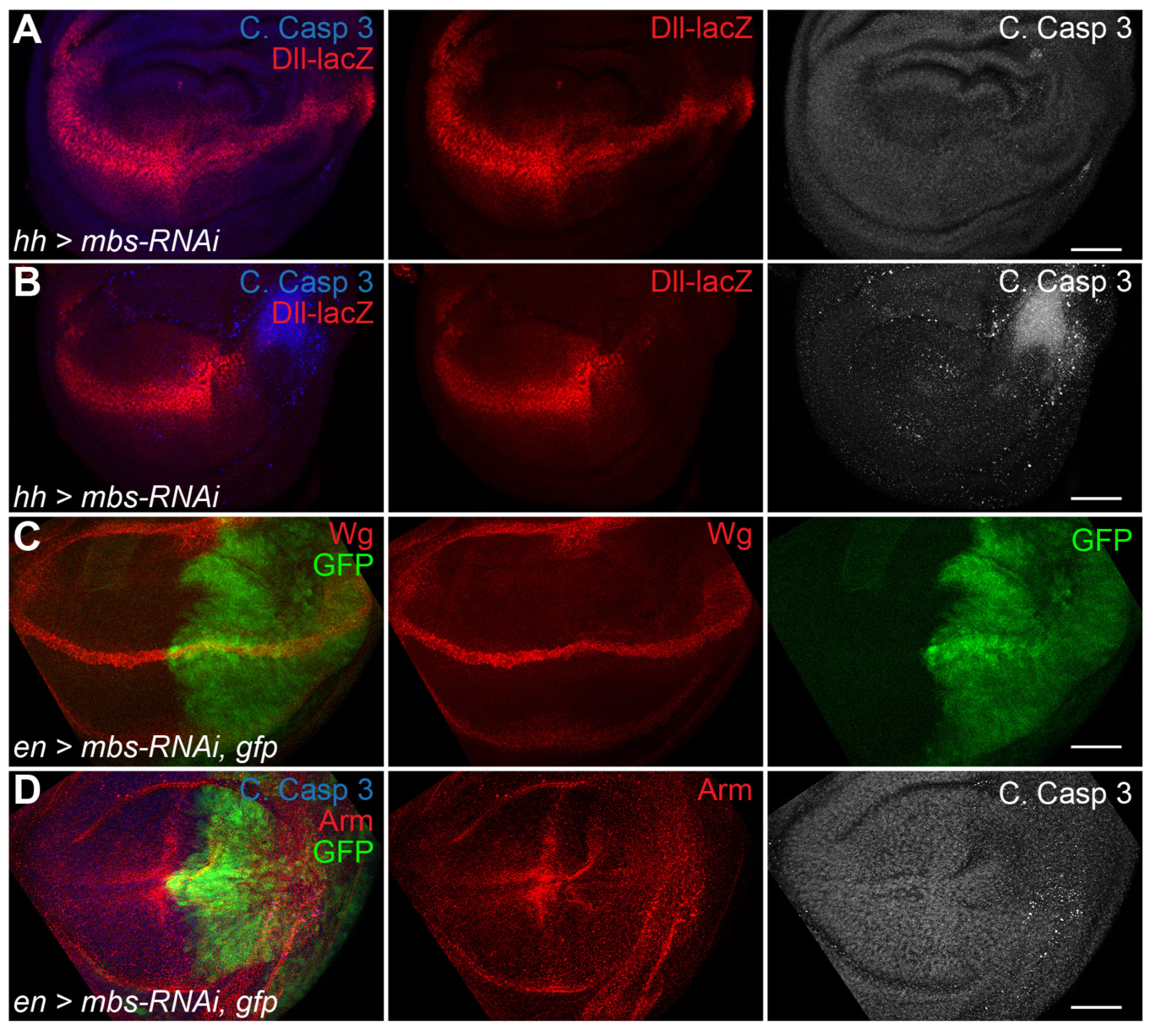
Myosin Binding Subunit, inhibits Wg activity but can induce cell death. Related to Figure 1. (A,B) *hh-Gal4* expressing *mbs-RNAi* in the posterior domain of the wing imaginal disc, reduces *Dll-lacZ* expression and can induce C. Casp 3 sporadically. (C) *en-Gal4* expressing GFP and *mbs-RNAi* in the posterior domain, does not affect Wg. (D) *en-Gal4* expressing GFP and *mbs-RNAi*, reduces stabilized Arm, without activating C. Casp 3. Scale bars: 50 μm.

**Figure S3.**
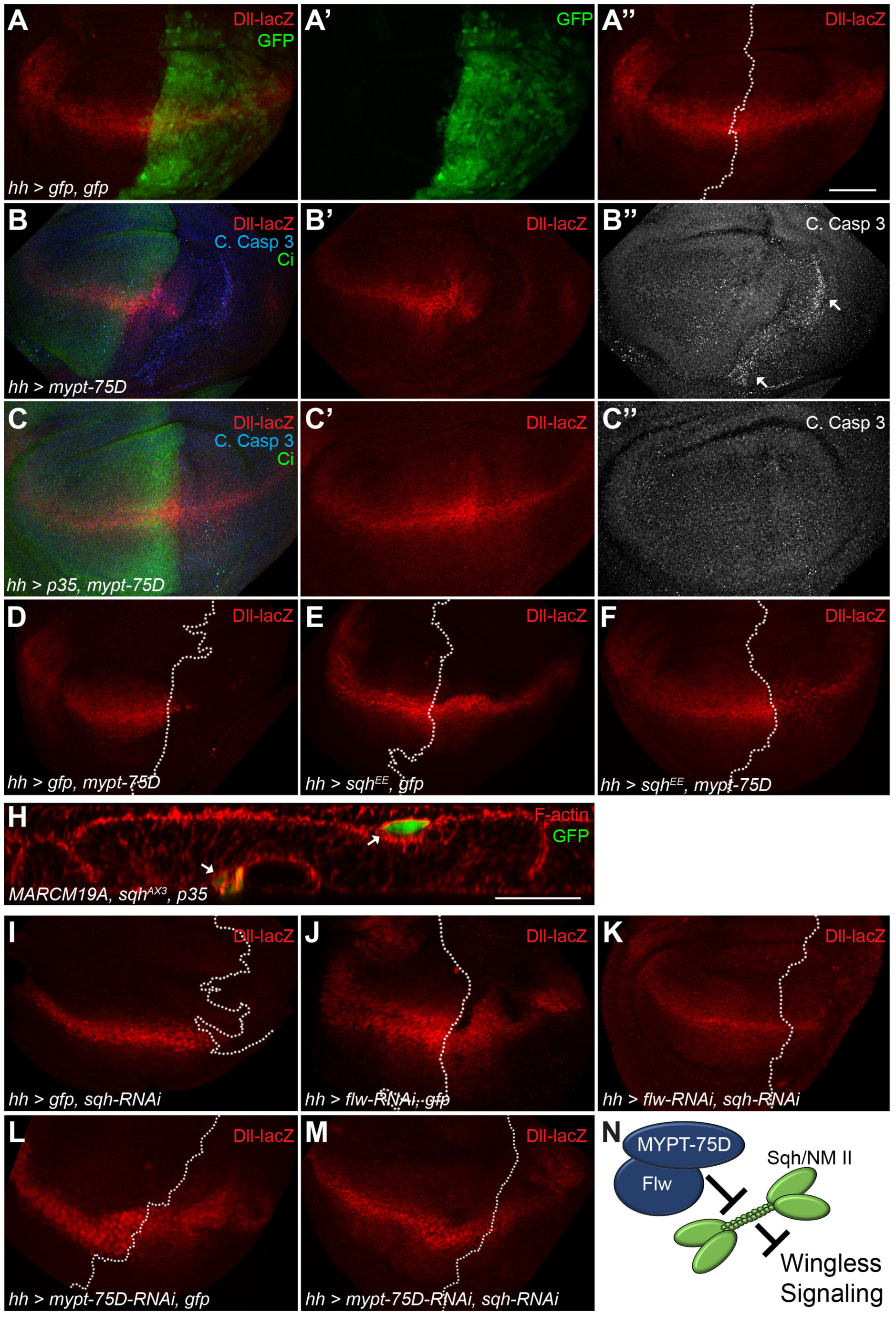
Myosin phosphatase mediates Wg activity and cell viability through NMII activation. Related to Figure 1. (A-A’”) Control wing disc with *hh-Gal4* in the posterior domain of the wing disc marked by GFP, stained for expression of *Dll-lacZ*. (A-A”). (B-C”) Wing discs stained for C. Casp 3 and Ci marking the anterior domain of the wing disc expressing MYPT-75D (B-B”), or P35 and MYPT-75D (C-C”). (D-F, I-M) Genetic interactions between myosin phosphatase components and Sqh utilizing *hh-Gal4* expressed in the posterior domain of the wing disc, stained for expression of *Dll-lacZ*. (D) GFP and MYPT-75D, (E) Sqh^EE^ and GFP, (F) Sqh^EE^ and MYPT-75D. (H) GFP-marked *sqh^AX3(null)^* MARCM clones expressing P35 (arrows mark cells extruding apically and basally). (I) GFP and *sqh-RNAi,* (J) *flw-RNAi* and GFP, (K) *flw-RNAi* and *sqh-RNAi*, (L) *mypt-75D-RNAi* and GFP, (M) *mypt-75D-RNAi* and *sqh-RNAi.* (N) Model for regulation of Wg signaling, myosin phosphatase regulates Wg activity through inhibition of NMII. Scale bars: (A-F, I-M) 50 μm, (H) 20 μm.

**Figure S4.**
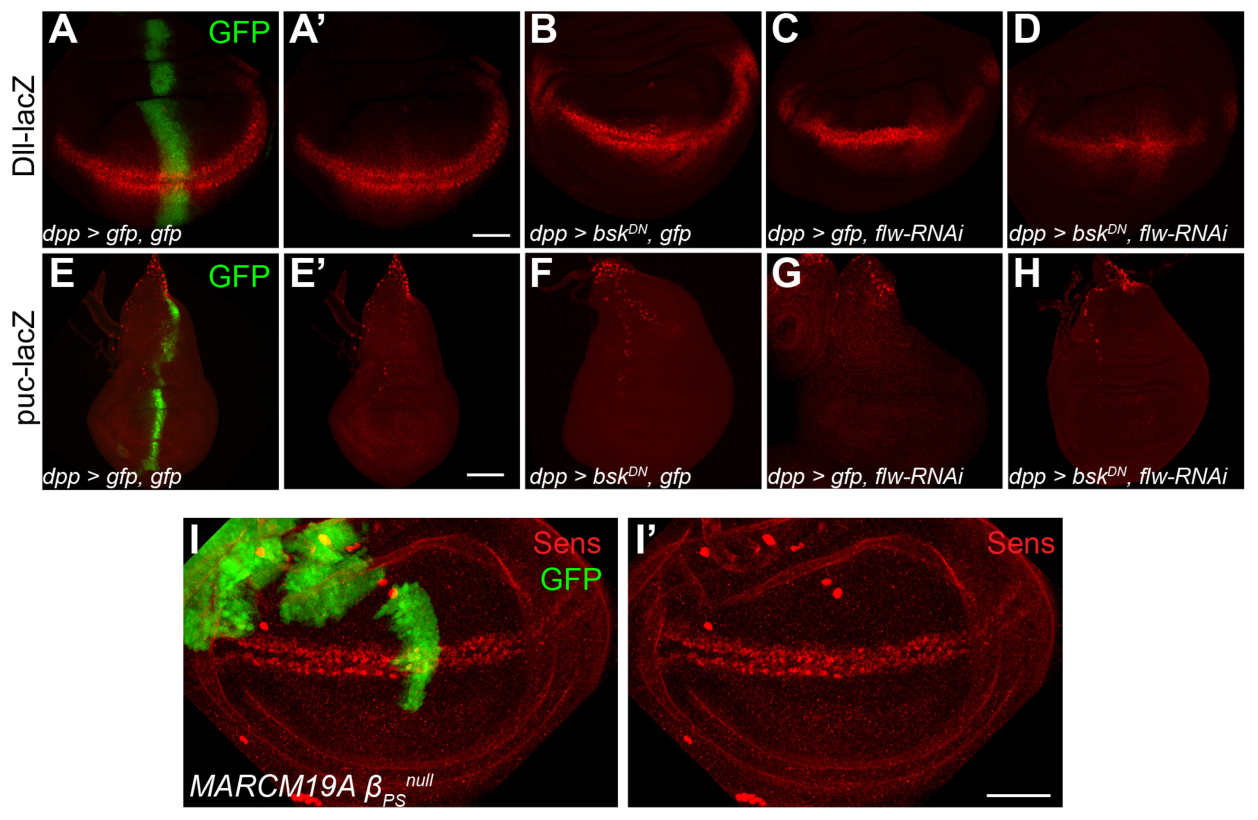
NMII does not regulate Wnt through JNK or integrin signaling in wing imaginal discs. Related to Figure 2. (A-D) *dpp-Gal4* expressing (A,A’) GFP, (B) Bsk^DN^ and GFP, (C) GFP and *flw-RNAi*, (D) Bsk^DN^ and *flw-RNAi*, stained for expression of *Dll-lacZ*. (E-H) *dpp-Gal4* expressing (E,E’) GFP, (F) Bsk^DN^ and GFP, (G) GFP and *flw-RNAi*, (H) Bsk^DN^ and *flw-RNAi*, stained for expression of *Dll-lacZ*. (I,I’) GFP-marked *β_PS_^null^* MARCM clones, stained for Sens. Scale bars: (A-D,I,I’) 50 μm, (E-H) 100 μm.

**Figure S5.**
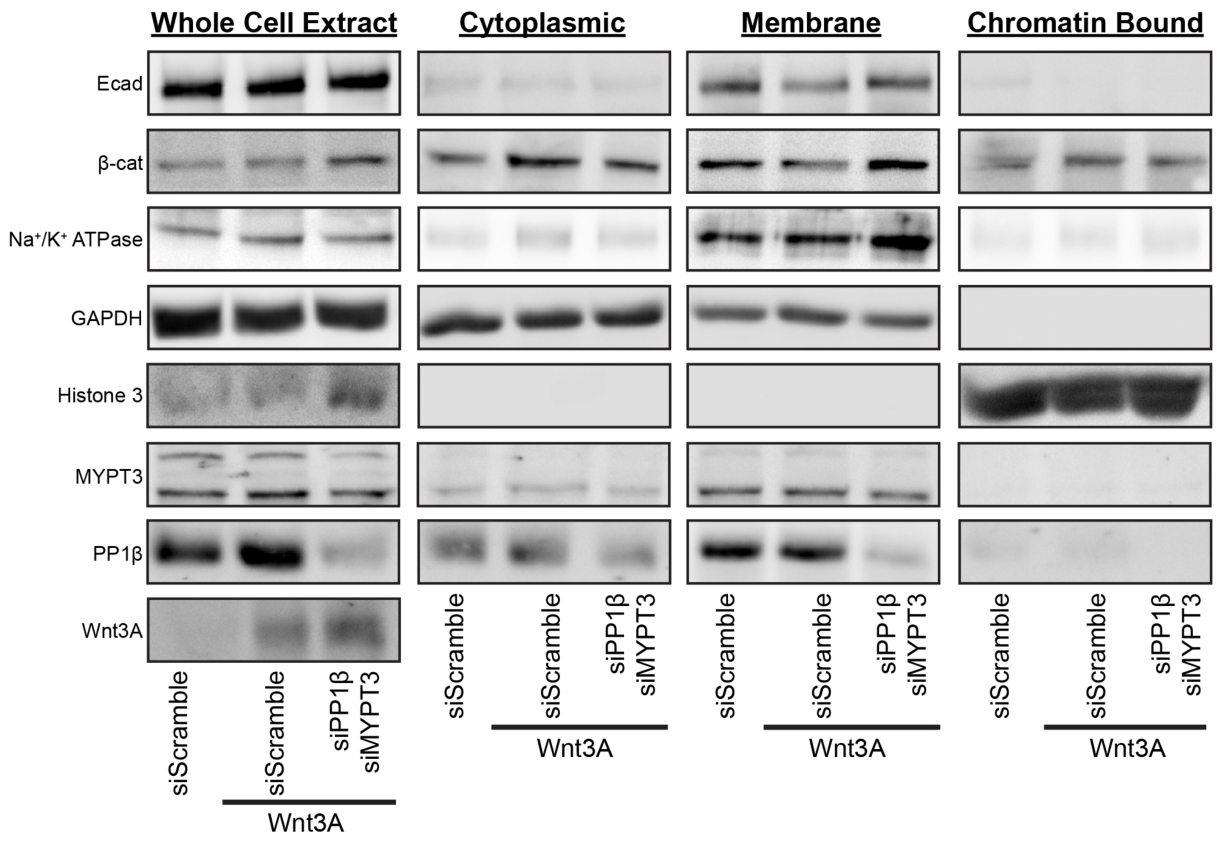
Complete fractionation of MCF7. Related to Figure 6. MCF7 cells transfected with or without Wnt3A, siPP1β and siMYPT3. Whole cell extracts and cell fractions were analyzed by western blots, probed for E-cad, β-cat, MYPT3, PP1β, and Wnt3A. GAPDH (whole cell extracts, and cytoplasmic), Na^+^/K^+^ ATPase (Membrane), and Histone 3 (Chromatin Bound) were used as loading controls for individual fractions. Blots shown represent one of 4 biological replicates.

**Figure S6.**
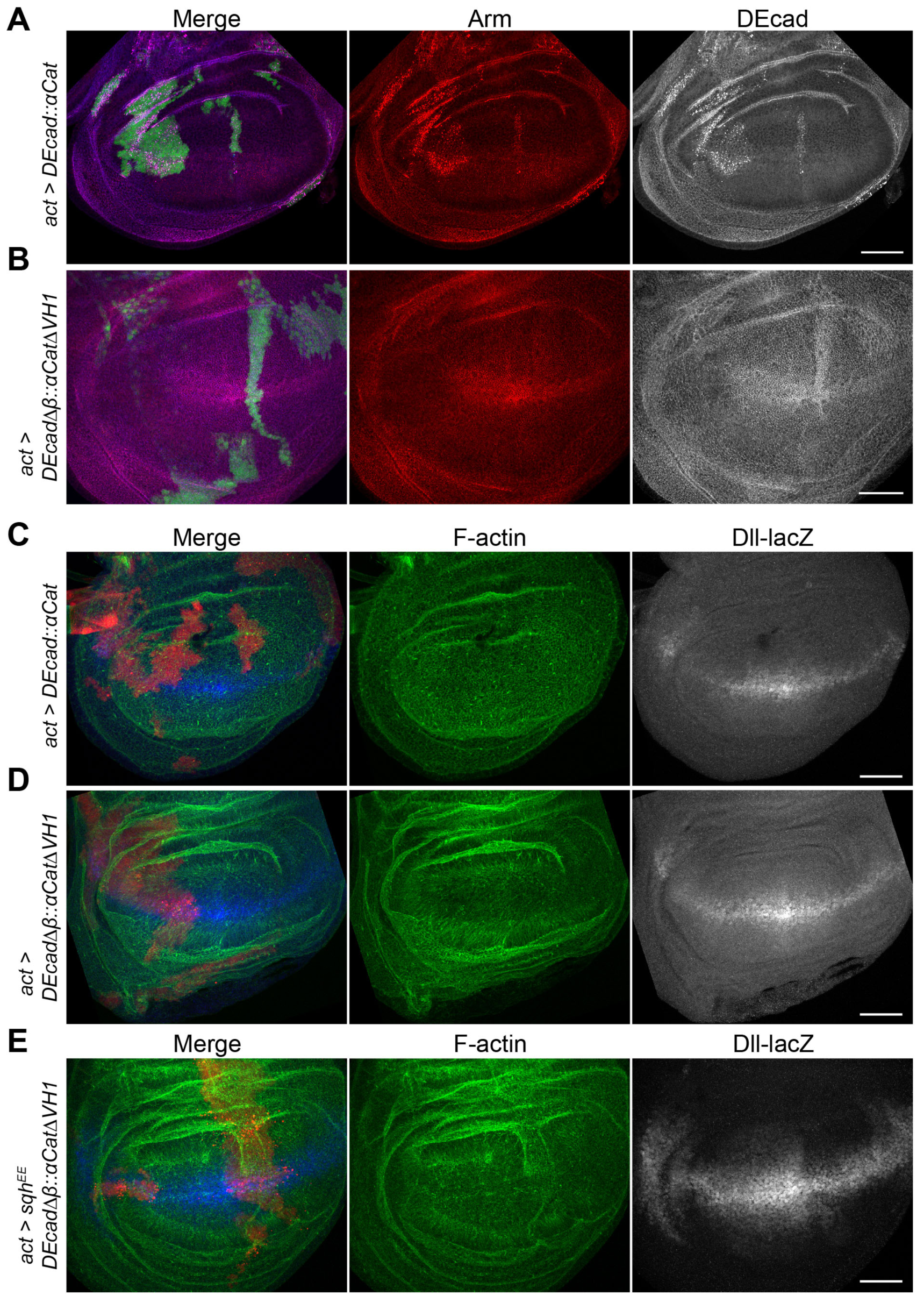
Full length and truncated DE-cad::α-cat fusion proteins effects on the wing imaginal disc. Related to Figure 7. (A,B) GFP-marked actin flip-out clones expressing (A) *DE-cad::α-cat* and (B) *DE-cad*Δ*β::a-catΔVHJ,* stained for Arm and DE-cad. (C-E) RFP-marked actin flip-out clones expressing (C) *DE-cad::α-cat* and (D) *DE-cad*Δ*β::a-catΔVH1*, and (E) *sqn*^EE^ with *DE-cad Apr. a-catAVHI,* stained for F-actin and *Dll-lacZ* expression. Scale bars: 50 μm.

